# Conformational Changes in the Negative Arm of the Circadian Clock Correlate with Dynamic Interactomes Involved in Post-transcriptionally Regulated Processes

**DOI:** 10.1101/2021.11.20.469315

**Authors:** Jacqueline F. Pelham, Alexander E. Mosier, Samuel C. Altshuler, Christopher L. Kirchhoff, William B. Fall, Lisa S. Baik, Joanna C. Chiu, Jennifer M. Hurley

**Author notes:** Authors contributed equally (Jacqueline F. Pelham, Alexander E. Mosier).

## Abstract

The circadian clock employs a transcriptional/translational negative feedback loop (TTFL) to anticipate environmental changes due to the Earth’s diurnal cycle, with regulation of organismal physiology believed to stem from temporal transcriptional activation by the positive arm. However, up to 80% of oscillating proteins do not have rhythmic mRNA, establishing circadian post-transcriptional regulation through unknown mechanisms. Given the pervasive conservation of the intrinsically disordered nature of negative-arm clock proteins, we hypothesized that post-transcriptional regulation may stem from conformational shifts in negative-arm proteins that time vacillations in the constituents of negative-arm macromolecular complexes to time cellular physiology. Our investigation of the negative arm clock protein in *Neurospora crassa*, FREQUENCY (FRQ), demonstrated temporal conformational fluidity correlated with daily changes in physiologically diverse macromolecular complex components. A parallel investigation of the macromolecular complexes centered around *Drosophila melanogaster* PERIOD (dPER) and human PERIOD (hPER2) found a similar number and physiological diversity of interacting partners in higher eukaryotes. Short linear motifs (SLiMs) associated with the interactors localized to disordered and phosphorylated regions on the PERs and FRQ, with disordered interactors oscillating in the macromolecular complexes over circadian time. This oscillation correlated with oscillations in post-transcriptionally regulated proteins, suggesting the negative arm may tune cellular physiology and proteostasis post-transcriptionally via vacillations in the circadian negative-arm macromolecular protein complexes.

## INTRODUCTION

Most organisms subjected to the Earth’s daily light/dark cycle have evolved a molecular mechanism to anticipate changes associated with this persistent oscillation in environmental conditions. This mechanism, called the circadian clock, increases survival, reproduction, and growth rates in organisms that maintain a clock ^1-3^. At the molecular level, the circadian clock comprises a transcription/translation negative feedback loop (TTFL) ^4,5^. The circadian cycle begins when the positive arm of the TTFL transcriptionally activates the negative arm, which in turn acts back on the positive arm to suppress this transcriptional activation, closing the loop. The negative arm then undergoes a tightly regulated half-life, allowing the positive arm to reactivate and begin the cycle anew ^4-8^. To influence physiology, the positive arm regulates gene promoters beyond the negative arm, controlling the rhythmic transcription of a large portion of the genome ^9-11^. For this reason, research has predominantly focused on time specific transcriptional activation by the positive arm to identify circadian output ^4,12,13^. However, recent research, including work in *Neurospora crassa (N. crassa), Drosophila melanogaster (D. melanogaster)*, and humans, has demonstrated that many proteins show circadian oscillations while their corresponding mRNA do not, though the source of this regulation is currently debated ^9,14-20^.

Beyond negative arm repression of the positive arm, the role of the negative arm in the regulation of output, if any, is not clear. What is known is that the negative arm forms a multitude of macromolecular complexes with a variety of proteins, including many that do not play a direct role in timekeeping ^21-24^. This paradigm is true in the TTFL of fungal, insect, and mammalian cells, where the core negative arm proteins, FREQUENCY in fungi (FRQ) and the PERIODs in animals (PERs) have been demonstrated to have a wide array of interactors, only some of which impact core clock functions ^21,23,25-28^. The conservation of large interactomes in negative-arm clock proteins is not surprising as negative-arm proteins from eukaryotic clocks are largely intrinsically disordered proteins (IDPs) ^6,29-33^. IDPs are a class of proteins known for their dynamic and heterogeneous ensemble of conformations whose inherent flexibility regulates binding to more extensive and transient interactomes as compared to globular proteins ^34-38^. Due to promiscuity in binding and demonstrated disorder, we hypothesized that negative-arm proteins possess temporal-specific conformational ensembles that time protein-protein interactions, allowing the negative arm to act as a temporal “Hub” protein to impart post-transcriptional regulation of output ^39-41^.

To investigate this hypothesis, we surveyed FRQ from *N. crassa* in its micro-environment over circadian time with the CRAFTY approach, using altered protease accessibility as a proxy to ascertain that the conformations of FRQ temporally oscillated ^30^. We coupled this with an investigation of the protein complexes centered around FRQ over the circadian day, demonstrating that the interactors correlated with conformational changes in FRQ and localized to disordered regions in the protein. To extend this paradigm to the negative arm in higher eukaryotes, we surveyed the complexes centered around the functional analog of FRQ in *D. melanogaster* (dPER), and humans (hPER2) ^21^, demonstrating that complexes centered around dPER are spatiotemporally specific and correlate with disordered regions. Among these negative arm interactors, there was an enrichment for proteins that had regions of intrinsic disorder, with temporal-specific disorder-to-disorder interactions occurring at the nadir of post-transcriptionally-regulated circadianly-oscillating proteins, signifying that negative arm proteins may play a role in proteostasis to synchronize the cell post-transcriptionally. Moreover, interactors played a variety of regulatory roles in the cell, suggesting potential pathways of post-transcriptional regulation beyond proteostasis.

## RESULTS

### CRAFTY Analysis Demonstrates Oscillations in the FRQ Conform-ome

IDPs are difficult to characterize using typical structural approaches due to heterogeneity in conformations and instability during over expression. Therefore, to demonstrate temporal changes in the ensemble of conformations (referred to here as the “conform-ome”) of the negative arm, we applied the CiRcadian nAtive FasT parallel proteolYsis (CRAFTY) approach to FRQ ^30^. CRAFTY analyzes the average changes in the conformations of a protein in a “native” lysate over circadian time to determine protein conformation, utilizing digestion/protease accessibility as a proxy for the protein microenvironment ^30,36^. We performed CRAFTY on age-matched *N. crassa* mycelial mats from a strain where the C-terminal end of FRQ was fused to a V5-10His-3FLAG tag inserted at the cyclosporin locus of an FRQ KO strain (τ=21.3+/-0.39 hrs) (strain WT8-1) (Supplemental Figure 1A) (see methods and materials). Mats were shifted to constant darkness (DD) before being sampled every 4hrs for 24 hours in triplicate over circadian time (Supplemental Figure 1B). Whole-cell lysates were processed through CRAFTY in parallel, analyzed by western blot, and quantified to determine the degradation rate of FRQ (Supplemental Figure 1C) (see methods). Our analysis found the rate of FRQ digestion varied over circadian time, with the most protease resistance at CT13.6 (∼155 sec) and the least at CT9.3 and CT22.1 (∼60 sec) (Figure 1A gray and Supplemental Figure 1C). This shift from a closed to an open conformation correlated with the progressive phosphorylation of FRQ, though increasing phosphorylation did not necessarily represent increasing protein accessibility at all time points, as has been reported previously ^42^.

**Figure 1.**
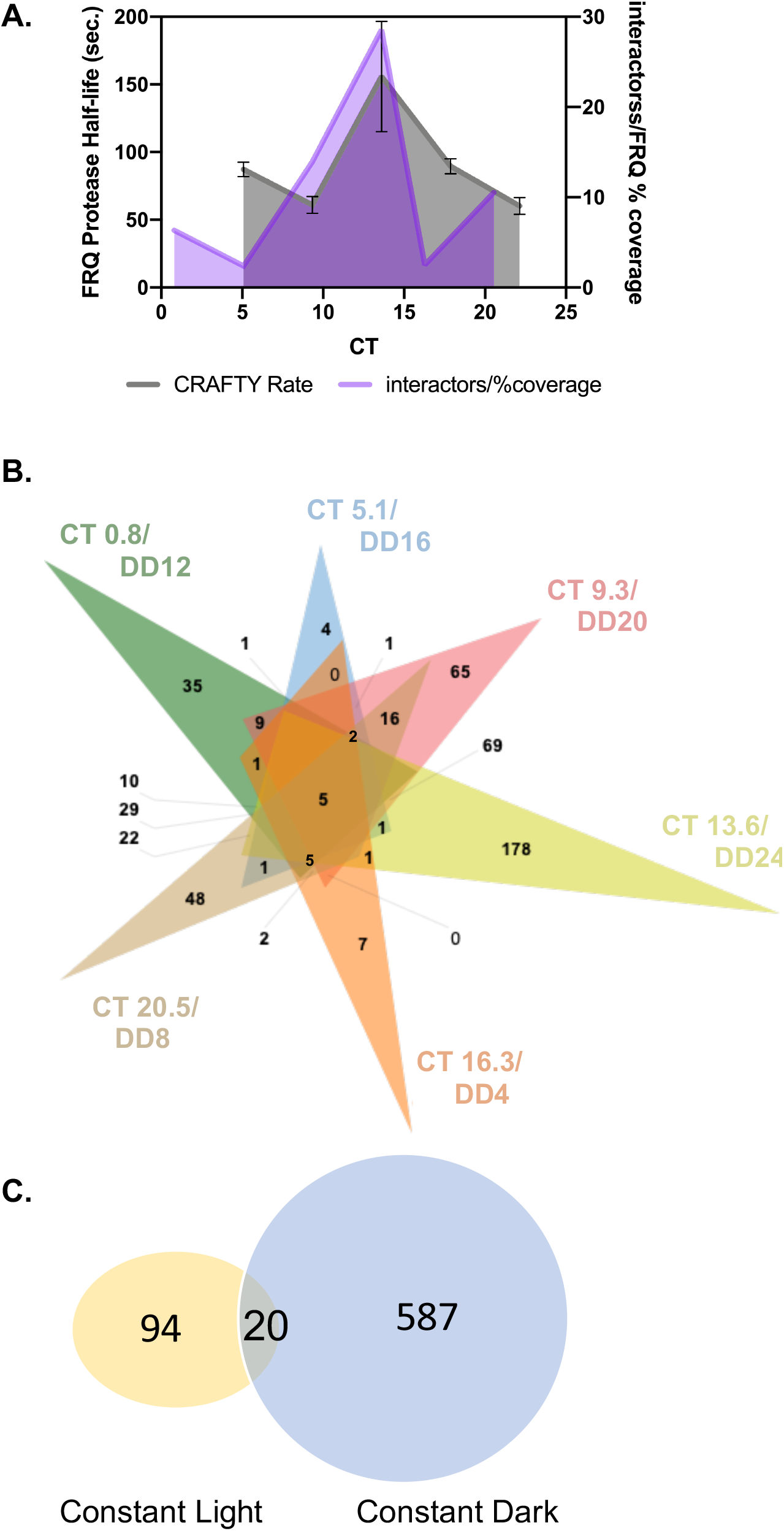
FRQ protease resistance and interactomes vary over the circadian day. A.) The protease resistance of FRQ, represented as protein half-life (grey), was plotted on the same graph as the % coverage normalized number of FRQ interactors (purple) at the same CT and shaded for ease of comparison. B.) Complex venn diagram displaying the overlap between proteins identified in complex with FRQ at different time points over the circadian day (CT0.8/DD12 in green, CT5.1/DD16 in cyan, CT9.3/DD20 in pink, CT13.6/DD24 in yellow, CT16.3/DD4 in orange, and CT20.5/DD8 in tan) ^99^. C.) Venn diagram displaying the overlap between proteins identified in complex with FRQ in LL (yellow) or DD (blue).

### Macromolecular Complexes Centered around FRQ Vacillate in Concordance with Changes in the FRQ Conform-ome

We hypothesized that the changes in the conform-ome of FRQ could be related to qualitative and quantitative changes in the proteins in FRQ macromolecular complexes. To validate this hypothesis, we utilized a strain with a V5-10His-3FLAG epitope-tagged FRQ allele inserted in the native FRQ locus (strain 1500-1) (τ=24.9+/-3.3hrs) (Supplemental Figure 2A, C and D). Using both the 1500-1 strain and a wild-type strain as a control (Ku70a), *N. crassa* was sampled over circadian time, with samples taken from constant light (LL), 4, 8, 12, 16, 20, and 24 hours after the light to dark transition (DD) (Supplemental Figure 2B). Macromolecular complexes centered on FRQ were purified from each time point via a two-step Ni^+^ agarose/αFLAG co-immunoprecipitation process and validated to bind to known FRQ-interacting partners WC-1, WC-2, and FRH (Supplemental Figures 2C and 2D). Co-immunoprecipitated proteins were analyzed using Nano-Spray Liquid Chromatography-Mass Spectrometry/Mass Spectrometry (NS-LC-MS/MS) (see methods) (Supplemental Dataset 1). The standard for detection from the NS-LC-MS/MS results was based on an empirical analysis of known FRQ-interacting proteins, and proteins were only included in the dataset if they had ≥5% total protein coverage or ≥4 unique peptides at a given time point in Scaffold. We further refined this list by eliminating proteins that were detected in less than two of the three triplicates at a given time point in the 1500-1 strain or detected in any Ku-70 time points (Supplemental Figure 2E).

As expected, FRQ was identified in all samples from the 1500-1 strain (Supplemental Figure 3A) and known FRQ-interacting partners FRH, CK-1, WC-1, and WC-2 were detected throughout the circadian day (Supplemental Dataset 2). 681 FRQ protein-protein interactions were identified, 658 of which were novel FRQ protein-protein interactions (Supplemental Dataset 1, compared to ^22^. Only three proteins beyond the FFC were found at all time points and eight additional interactors were found across four or more time points (Supplemental Figure 3A and Supplemental Dataset 2). Conversely, 433 proteins were unique to a single circadian time point, with the number of interacting partners varying from as few as 22 interactors at CT5.07/DD16 to as many as 357 interactors at CT13.6/DD24 (Figure 1B, Supplemental Dataset 1, and Supplemental Dataset 2). When we compared the peak of protease resistance via our CRAFTY analysis to the peak of protein-protein interactions with FRQ via our NS-LC-MS/MS analysis, we found peak interaction aligned with peak protease resistance (Figure 1A).

We next compared the FRQ centered-macromolecular complexes from LL to the FRQ centered-macromolecular complexes from DD (Supplemental Dataset 1). We found that while 20 proteins were involved in FRQ macromolecular complexes in both LL and DD samples, 94 proteins were specific to LL FRQ macromolecular complexes, and 567 proteins were specific to DD FRQ macromolecular complexes (Figure 1C). This data supports previously published work demonstrating a difference between LL and DD FRQ ^22,43,44^.

### Predicted Short Linear Motifs in FRQ-centered Macromolecular Complexes Cluster in Disordered Regions

IDPs are known to facilitate binding through conserved Short Linear Motifs (SLiMs), regions less than 15 residues in length that are characterized by residue-pattern conservation in regions with otherwise low levels of explicit sequence conservation ^45-47^. Post-translational modifications (PTMs) commonly regulate SLiMs ^48,49^ and FRQ has over 100 identified phosphosites, mainly in the disordered regions of FRQ (Figure 2A and B) ^22,30,31,48-52^. To determine if there was a correlation between SLiMs, disorder, and phosphorylation in FRQ, we computationally investigated the molecular grammar of FRQ for SLiMs using a eukaryotic linear motif (ELM) predictor with a taxonomic context filter specific for the fungal kingdom and a motif probability cutoff of 100 ^53^. We then manually curated the identified SLiMs by removing any for which a known SLiM binder was not identified in the FRQ interactome. The remaining SLiMs were mapped to a linear representation of FRQ in addition to the known phosphorylation sites from Baker et al. 2009^22^ (Figure 2B and Supplemental Table 1). Thirteen FRQ-interactor-specific SLiMs were identified, color-coded by class, and mapped to their position (or positions if identified in multiple locations) on FRQ (Figure 2B and Supplemental Table 1). We then plotted the predicted regions of disorder and disordered to order transition in FRQ using IUPred2A(L) and ANCHOR2 respectively (Figure 2A). We found that SLiMs for validated FRQ interactors were ubiquitously found in regions of disorder, with a high tendency of disordered to ordered transition, and in regions that were near or at validated phosphorylation sites (Figure 2).

**Figure 2.**
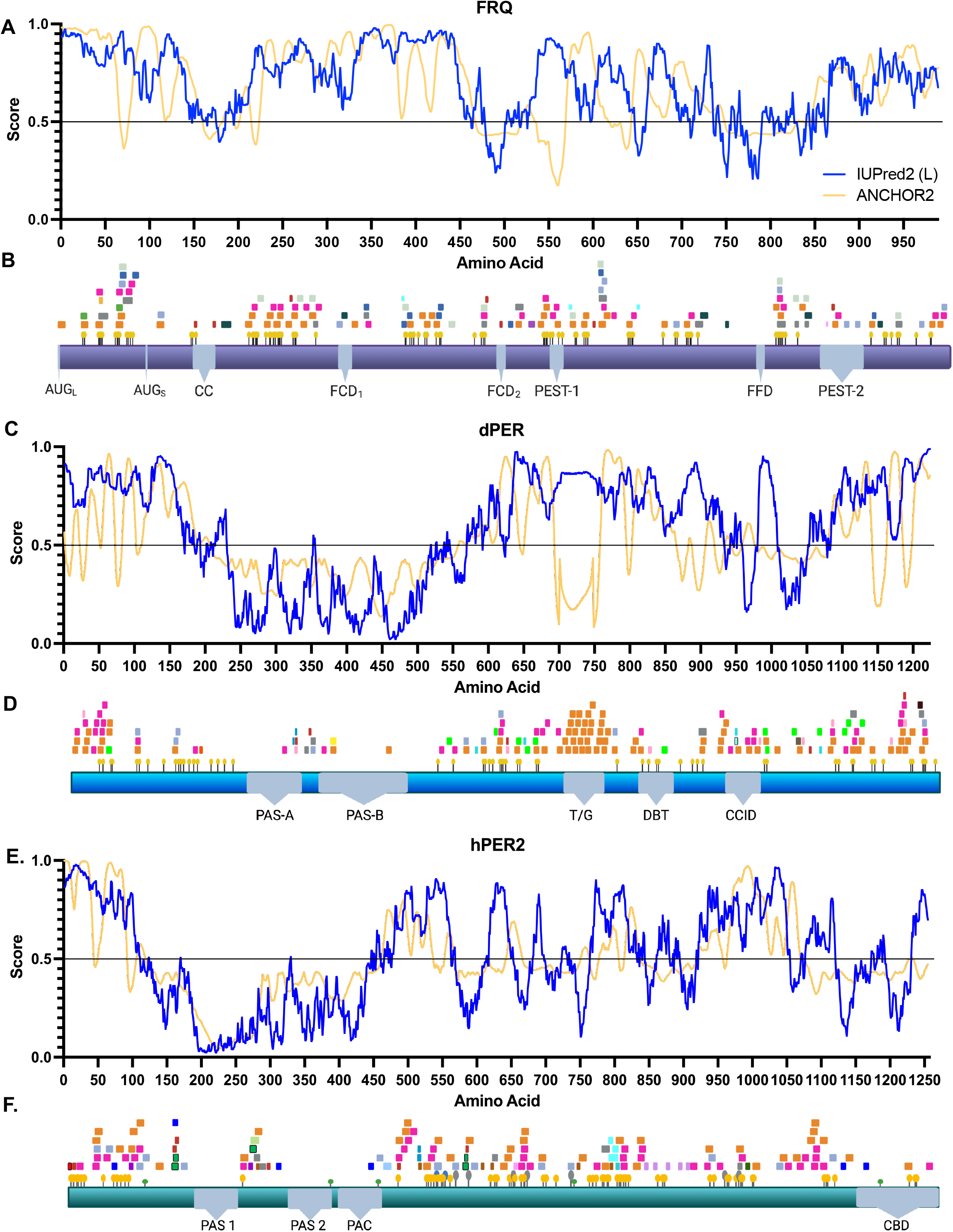
Sequential analysis of negative arm elements demonstrates interactome-confirmed SLiM regulation sites in disordered regions. A., C., and E.) A linear representation of the disorder propensity and disorder to order transition upon binding for FRQ, dPER, and hPER2 respectively, calculated using IUPred2A(L) and ANCHOR2 ^89^. B., D., and F.) A linear map of ELM computationally-predicted FRQ, dPER, and hPER2 SLiMs respectively that had interactors identified in the LC-MS/MS analysis ^53^. SLiMs and PTMs ^22^ were mapped to their location on FRQ, dPER, and hPER2 (see legend in Supplemental Table 1). Known FRQ domains highlighted in grey and phosphosites as yellow pins. Long and short start codons: AUG_L_ and AUG_S_, CC: Coiled coil domain for FRQ dimerization, FCD_1_ and FCD_2_: FRQ and CK1 interaction region, FFD: FRQ and FRH interaction region ^30^. Known dPER domains highlighted in grey PAS-A and PAS-B: PER-ARNT-SIM domains, T/G: Threonine glycine rich region, DBT: DOUBLETIME interacting region, CCID: dCLK and CYC inhibition domain) and phophosites as yellow pins. Known hPER domains highlighted in grey (PAS-1 and PAS-2: PER-ARNT-SIM domains, PAC: C-terminal PAS domain, CBD: Period C Binding domain), phophosites as yellow pins, glycosylation as grey pins, acetylation as blue pins, and ubiquitination as green pins ^100^.

### Macromolecular Complexes in higher eukaryotes are spatiotemporally specific, regulated by phosphorylation, and centered in disordered regions

To determine if the correlation between SLiMs and regions of disorder was conserved in higher eukaryotes, we performed affinity purification coupled with MS/MS to survey the macromolecular complexes centered around *Drosophila* PERIOD (dPER) in a 12h light and 12hr dark (12:12 LD) entrainment scheme. To do so, dPER was immunopurified at ZT 3, 16, 20, and 24 via a 3xFLAG antibody from cytosolic and nuclear fractions from *D. melanogaster* heads in an N-terminal, 3xFLAG-tagged, dPER line with an untagged line as a control. Samples were then analyzed by MS/MS and proteins with ≥4 unique peptides in at least 2 out of 3 replicates in the 3xFLAG-tagged dPER, but not the untagged line, were considered valid dPER interactions. As expected, we found TIMELESS (TIM) and DOUBLETIME (DBT) in all timepoints (Supplemental Dataset 3). In addition to TIM and DBT, 248 unique proteins were detected in complex with dPER (Figure 3A and B, and Supplemental Dataset 3), but only three of these were found in complex with dPER at all time points, suggesting that like FRQ, dPER has an extensive and temporally-regulated interactome. Of the 248 interactions, 89 and 115 proteins were unique to the nucleus and cytosol respectively (Figure 3B and Supplemental Dataset 3).

**Figure 3.**
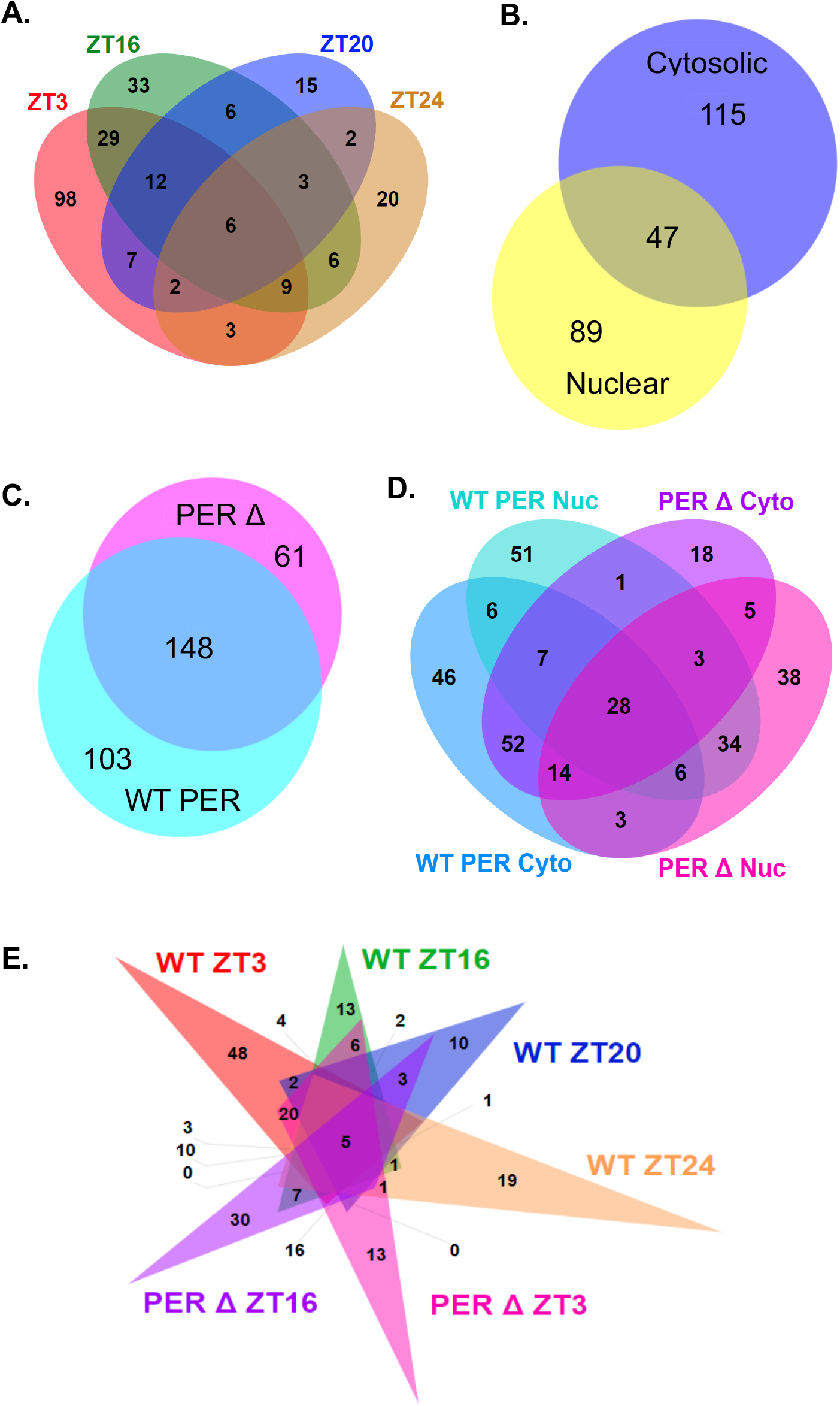
Macromolecular complexes centered around dPER are spatiotemporally specific. A.) Comparison of proteins identified in complex with dPER in WT samples grouped by timepoint (ZT3-Salmon, ZT16-Green, ZT20-Blue, and ZT24-Tan. B.) Comparison of proteins identified in complex with dPER in WT samples grouped by subcellular localization (Cytosolic-Blue and Nuclear-Yellow). C.) Comparison of dPER interactors in all WT and PERΔ samples (WT-Cyan and PERΔ -Pink). D.) Comparison of WT and PERΔ grouped by subcellular localization (WT/Cytosolic-Blue, WT/Nuclear-Cyan, PERΔ/Cytosolic-Purple, PERΔ/Nuclear-Pink). (E.) Comparison of WT and PERΔ grouped by timepoint (WT/ZT3-Salmon, WT/ZT16-Green, WT/ZT20-Blue, WT/ZT24-Tan, PERΔ/ZT3-Pink, PERΔ/ZT16-Purple).

We next computationally investigated the molecular grammar of dPER for the tendency for disorder, disordered to order transition, and SLiMs using IUPred2A(L), ANCHOR2, and a eukaryotic linear motif (ELM) predictor as above, identifying 14 SLiMs (Figure 2C and D and Supplemental Table 1). Moreover, we similarly surveyed the molecular grammar of hPER2 based on interactors from a previously published study, identifying 18 SLiMs (Figure 2E and F and Supplemental Table 1) ^21^. In both cases, the majority of SLiMs mapped to regions with a higher tendency for protein disorder and phosphorylation, suggesting that the relationship between disorder, phosphorylation, and binding was conserved in animal clocks (Figure 2).

To investigate the effect of phosphorylation of negative arm proteins on their interactomes, we repeated our interactome analysis on a *Drosophila* strain with an internal deletion in the DBT binding domain on dPER (755-809) (dPERΔ) that results in strain arrhythmicity and altered phosphorylation (see methods) ^54^. 3xFLAG-tagged dPER and dPERΔ had 148 interactors in common, 103 and 61 interactors unique to dPER and dPERΔ respectively (Figure 3C). There were further differences between dPER and dPERΔ when comparing spatiotemporal interactions, suggesting an important role for DBT and phosphorylation in both the location and timing of complex constituents (Figure 3D, 3E, and Supplemental Dataset 3).

### Conserved Interaction Classes Suggest Diverse Regulatory Roles for Negative-Arm Complexes

Given the time-of-day specific macromolecular complexes that form in the negative arm, and that disordered scaffold proteins have been suggested to serve as “Hub” proteins, highly connected proteins that are essential in a protein-protein interaction network, we proposed the negative arm was ideally suited to act as a temporal “Hub” protein ^**55**,**56**^. To investigate the physiological effect the negative arm could impart post-transcriptionally by acting as a time-specific “Hub” protein, we surveyed the interactor/SLiM-class parings conserved across our datasets at the single SLiM level. One common SLiM class, the highly-conserved 14-3-3 proteins, which identify phosphorylated substrates in disordered regions to regulate cell division and other processes, was seen in all species (Figures 2B, D, and F, and Supplemental Table 1) ^57-65^. Correspondingly, 10 FRQ and 3 hPER2 and dPER 14-3-3 SLiMs resided in regions of disorder (Figure 2) ^66^. Of note, 14-3-3 proteins were not among the dPERΔ interactors, emphasizing the importance of phosphorylation in regulating the formation of negative-arm “Hub” complexes.

Consistent with the importance of phosphorylation, 21, 12, and 16 kinases were found in complex with FRQ, dPER, and hPER2 respectively, and phosphorylation SLiMs aligned with many of the experimentally-verified phosphorylation sites (Supplemental Datasets 1 and 3) ^22,50,67^. Since not all kinases found in negative-arm macromolecular complexes target the negative arm, these kinases may serve to phosphorylate other negative-arm binding proteins to affect cellular physiology ^22,50-52^. For example, Cyclin-dependent kinase Cell Division Control Protein 2 (CDC-28) in *N. crassa*, a homolog of human CDK-1, acts as a cell cycle gatekeeper and must be phosphorylated before the cell can transition beyond S- and M-phases ^68-70^. CDC-28 was identified as an FRQ interactor at CT20.53/DD8, the time known to correlate with the clock regulation of entry to S-phase and the same time as the zenith of several post-transcriptionally rhythmic S-phase associated proteins, (MCM3, MCM4, and ORC3) (Supplemental Dataset 1 and Supplemental Figure 4) ^71^.

To explore the resonating effects of temporal changes to the macromolecular complexes centered around FRQ and dPER at the interactome level, we created a program that utilizes the STRING database to generate a temporal protein-protein interaction network (PIN) based on our interactome analysis ^72^. Our PIN employed Gene Ontological analysis (GO) to assign interactor functions using the TopGo package adapted from our Multi-Omics Selection with Amplitude Independent Criteria (MOSAIC) program and PANTHER to assign GO terms to each primary interactor for both dPER and FRQ, with predicted secondary interactors for FRQ (Figure 4A and Supplemental Figure 5 and 6) ^73-76^. For ease of visualization, the program only reported GO terms if the GO term had at least 20 assigned proteins. The PIN maps displayed the different impact in biological effect that negative-arm macromolecular complexes may have at different circadian time points and in different cellular compartments (Figure 4A and Supplemental Figure 5 and 6). We also created word clouds to represent the magnitude of enrichment of each of the interactor GO categories for FRQ, dPER, and hPER2 (Figure 4B, C, and D).

**Figure 4.**
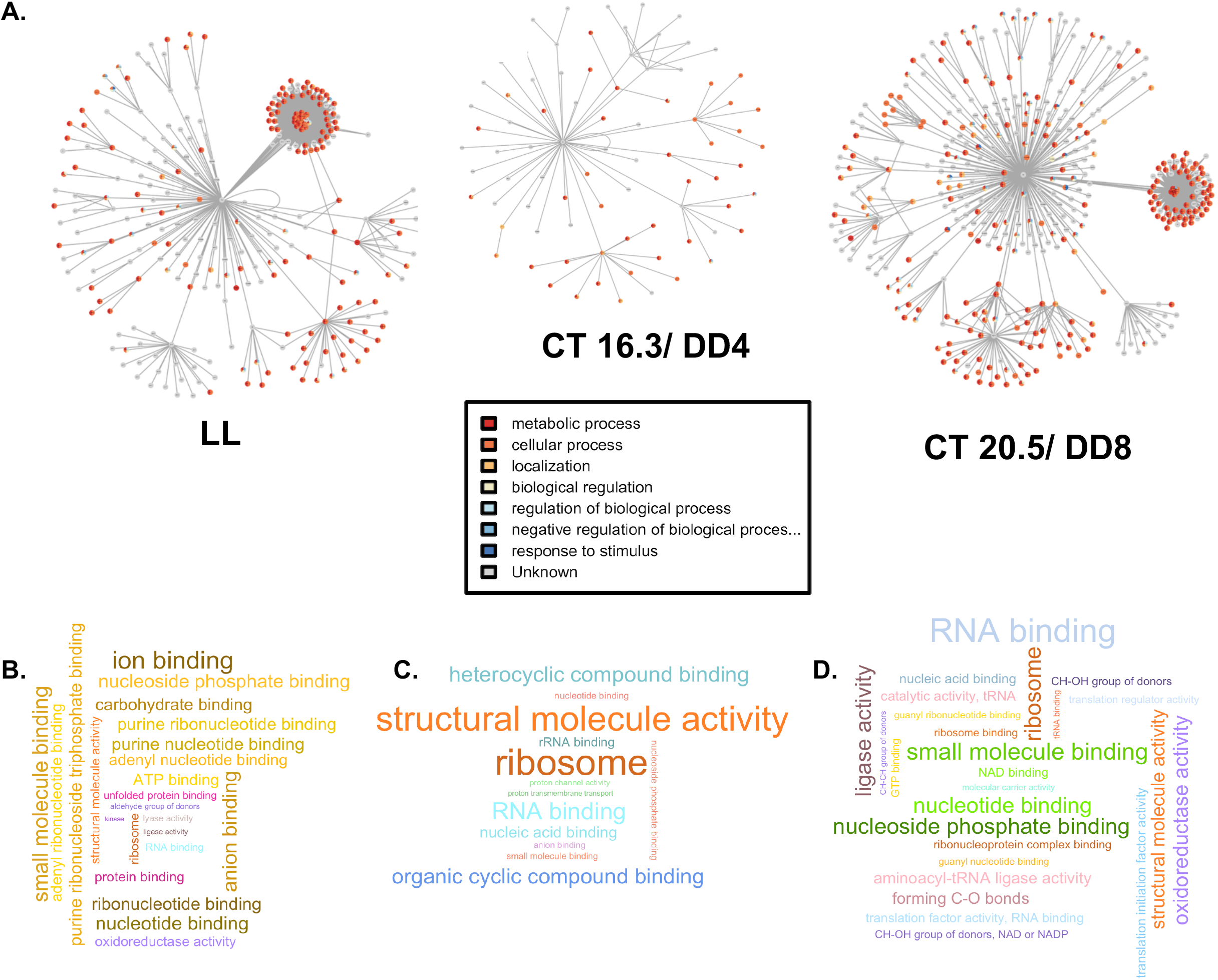
Negative-arm macromolecular complexes oscillate in composition and physiological function over the circadian day. A.) Protein interaction maps exhibiting the FRQ interactome at different timepoints (shown as examples are LL, CT16.3/DD4, and CT20.5/DD8). FRQ is in the center with radial lines connecting to proteins identified by LC-MS/MS. Predicted secondary interactors are connected to their respective primary interactor. Nodes are color coded for the secondary GO term shown. B.-D.) Word clouds related to the enriched gene ontologies of the interactions, created using terms with a higher than 2-fold enrichment and p-value >0.05, with the magnitude of each term represented by the -log10 of its p-value for FRQ (B.), dPER (C.), and hPER (D.) and colors indicating each GO family (Color Key in Supplemental File 2).

We found that metabolic GO terms accounted for the majority of interactors of FRQ between CT5.1 and CT13.6 but fell below 50% later in the circadian day (Figure 4A and Supplemental Dataset 2). The timing of the increase in metabolic proteins in FRQ macromolecular complexes correlated with known post-transcriptional regulation of metabolism in the circadian evening in *N. crassa* ^19^. Further, our analysis also showed an enrichment for RNA binding and structural constituents of the ribosome in *N. crassa, D. melanogaster*, and humans, including the 60S ribosomal protein L8 and several eukaryotic initiation factors that are found in all three datasets (e.g. dPER: RpL8, FBgn0261602, hPER2: RPL8, P62917, and FRQ: *crp-41*, NCU4779) (Figure 4B, C, and D, Supplemental Dataset 1, Supplemental Dataset 3, and Supplemental Dataset 2). This is significant as a cause of circadian influence on post-transcriptional regulation has been traced to translational rates ^19,20,77,19^.

### IDP/IDP Interactions May Impart Circadian Post-Transcriptional Regulation

IDPs are known to be stabilized through interactions with their binding partners and proteostasis is predicted to be circadianly regulated ^78-82^. Therefore, beyond the role of the negative arm as a “Hub” protein, another possible mechanism for circadian post-transcriptional regulation could be the regulation of the half-life of negative-arm interacting proteins. Though order- to disordered-region binding is more frequently described in the literature, recent data has shown that IDPs can also interact with other IDPs to affect molecular change ^83^. We hypothesized that negative-arm binding to IDPs could lead to post-transcriptional regulation of these IDPs via proteostasis and, if so, that IDPs would be overrepresented in the interactomes. To determine how pervasive these IDP/IDP interactions were, we examined the extent the proteins found in complex with the negative arm were disordered. Four disorder predictors (VLXT, VSL2b, IUPred2A(L), and PV2) were used to predict disorder in interactors and the average disorder score was used to classify each interactor as either an Intrinsically Disordered Protein (IDP, disordered score > 30%), a Protein with Disordered Regions (PDR, disordered score < 30% with 30 residue disordered ends or 40 residue disordered internal regions), or an Ordered Protein (OP, disordered score < 30% with no end or internal disordered regions), as was defined by Deiana et al. 2019 (Supplemental Dataset 4) ^84-90^.

In the case of FRQ, just under half (267 or 45.5%) of interactors were defined as IDPs by our classification (Supplemental Figure 3B). As we determined the *Neurospora* proteome was comprised of 54.5% IDPs, IDPs were significantly underrepresented in FRQ macromolecular complexes, contrary to our prediction (Fisher’s exact test p < 10^−5^) (Supplemental Figure 3B). Conversely, PDRs and OPs were overrepresented in the FRQ interactome (Fisher’s exact test p < 0.004) (Supplemental Figure 3B). We next examined the interactor disorder classification (IDP, PDR, and OP) by time of day of binding to FRQ. Contrary to the underrepresentation of IDPs in the overall dataset, IDPs were enriched in the late circadian night/early circadian morning, while IDPs/PDRs/OPs were more evenly distributed in the late circadian morning/early circadian afternoon, suggesting a circadian rhythm to the binding of IDPs (Figure 5A and Supplemental Dataset 5). To determine if the time-of-day specific increase in FRQ/IDP interactions could be explained by stoichiometric ratios or if these proteins may represent cases of circadian post-transcriptional regulation due to FRQ-induced stabilization, we identified interactors that were reported to oscillate with a circadian period at the protein level (Supplemental Dataset 5) ^19^. When we compared the interaction time to the zenith of the corresponding protein levels of FRQ interactors, we found that PDRs and OPs interacted with FRQ at their zenith, while IDPs predominantly interacted with FRQ during their nadir (Figure 5B, C and D). This suggests FRQ/IDP interactions are timed to occur at the IDPs nadir levels rather than occurring based on passive interaction due to stoichiometry.

**Figure 5.**
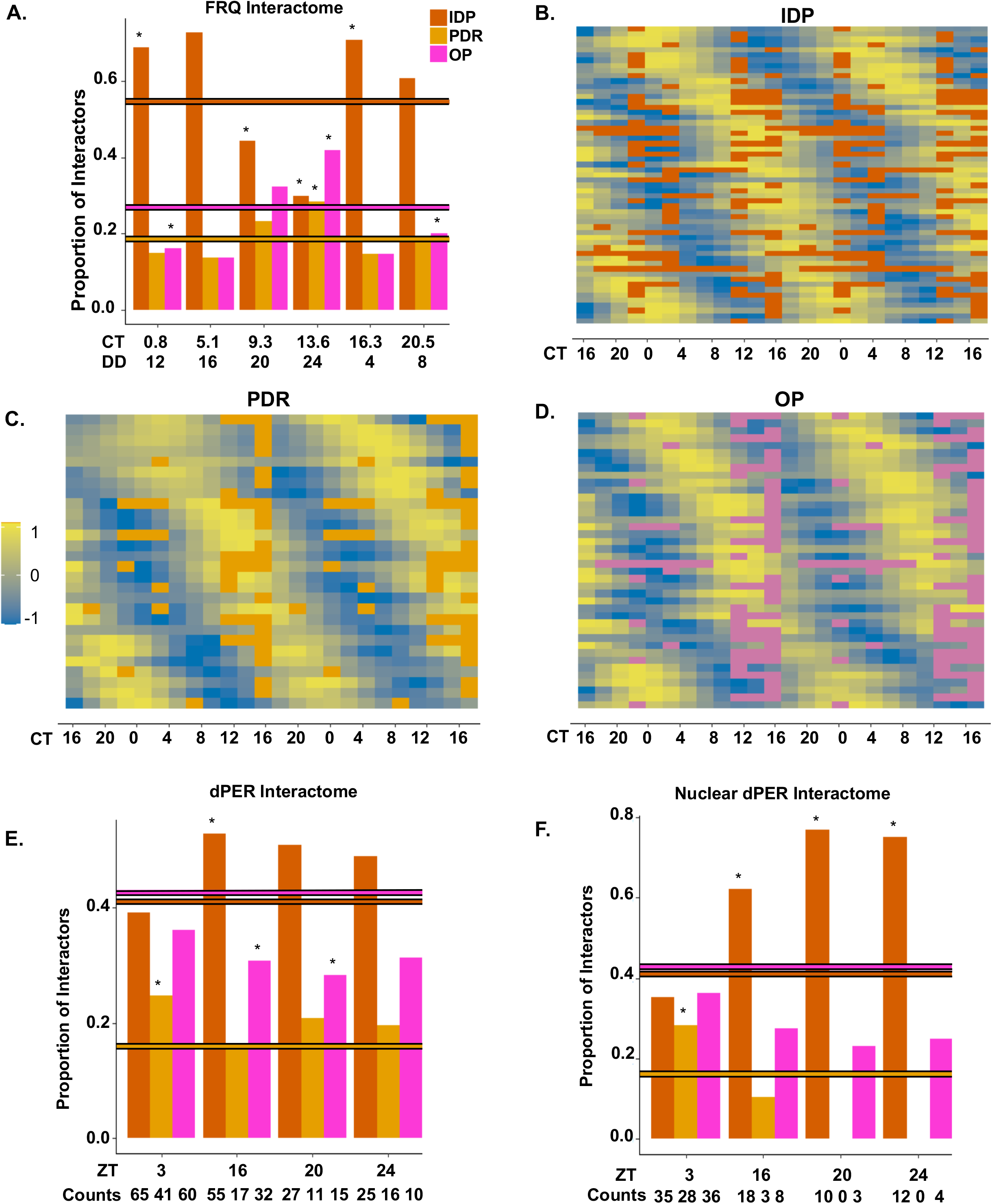
IDP/negative-arm protein interactions oscillate over the circadian day. A.) Bar plot showing the proportion and enrichment of IDPs, PDRs, and OPs in FRQ macromolecular complexes at a given time point as compared to the proportion of IDPs, PDRs, and OPs in the full *N. crassa* proteome (horizontal lines) (p-values assessed by Fisher’s exact test * indicating p<0.05). B.-D.) Heat maps showing global protein levels for oscillating, FRQ-interacting, IDPs (B.), PDRs (C.), and OPs (D.) as reported in Hurley et al. 2018 ^19^. These global protein levels are overlaid with the times of day that the corresponding protein interacted with FRQ for each IDP (orange), PDR (yellow), or OP (purple) ^19^. E. and F.) Bar plot showing the proportion and enrichment of IDPs, PDRs, and OPs in dPER macromolecular complexes at a given time point as compared to the proportion of IDPs, PDRs, and OPs in the full *D. melanogaster* proteome (horizontal lines) (p-values assessed by Fisher’s exact test p<0.05) in the total *D. melanogaster* interactome (E.) and the nuclear *D. melanogaster* interactome (F.).

Though we did not have temporal proteomic data to compare to, we performed similar analysis to quantify the extent of intrinsic disorder in dPER macromolecular complexes. Unlike FRQ, there was no enrichment or depletion of OPs, IDPs, and PDRs in the dPER interactome (Figure 5E and Supplemental Dataset 5). However, the nuclear dPER fraction displays a circadian pattern of interaction with IDPs, with a significant enrichment of IDPs at ZT 20 and 24 (Figure 5F and Supplemental Dataset 5).).

## DISCUSSION

A discrepancy between the circadian oscillations of mRNA and their cognate proteins has been established, though the mechanisms of this post-transcriptional regulation remain elusive ^14-16,19,20^. As we and others have shown that negative-arm proteins can modulate conformation over the circadian day, we hypothesized that the conserved intrinsic disorder documented among negative arm proteins could allow for conformational changes that correlate with changes in negative-arm protein-protein interactions to effect circadian post-transcriptional regulation ^29-31,42,91^. To validate this hypothesis, in this report we performed a CRAFTY analysis of the FRQ conformation over circadian time, a temporal analysis of the macromolecular complexes centered around FRQ and dPER using MS/MS, and mined data from previous analyses on hPER2 interactors ^21^. While the mechanistic relationship between shifts in the FRQ macromolecular complexes and the FRQ conform-ome remain unclear, the conformational vacillations of the negative-arm protein FRQ correlated with the peak numbers of interactors identified in our investigation (Figure 1). In concordance with this, the extent of proteins involved in negative-arm macromolecular complexes, the role of phosphorylation in dPER protein complexes, and the demonstration that SLiMs were in disordered and phosphorylated regions (Figures 1, 2, and 3), suggest that changes in phosphorylation states of negative-arm proteins allow for variations in the availability of negative-arm SLiMs and thereby the components of macromolecular complexes. Though our methods cannot determine if the identified interactions are direct, our computational analysis showed that many interactors had cognate SLiMs, providing some evidence that many of the proteins may directly interact with FRQ, dPER, and hPER. As many eukaryotic negative arm clock proteins are known to be IDPs, we propose that disorder in the negative arm proteins allows for conformational flexibility that times the formation of macromolecular protein complexes in all eukaryotic clocks (Figure 6) ^31^.

**Figure 6.**
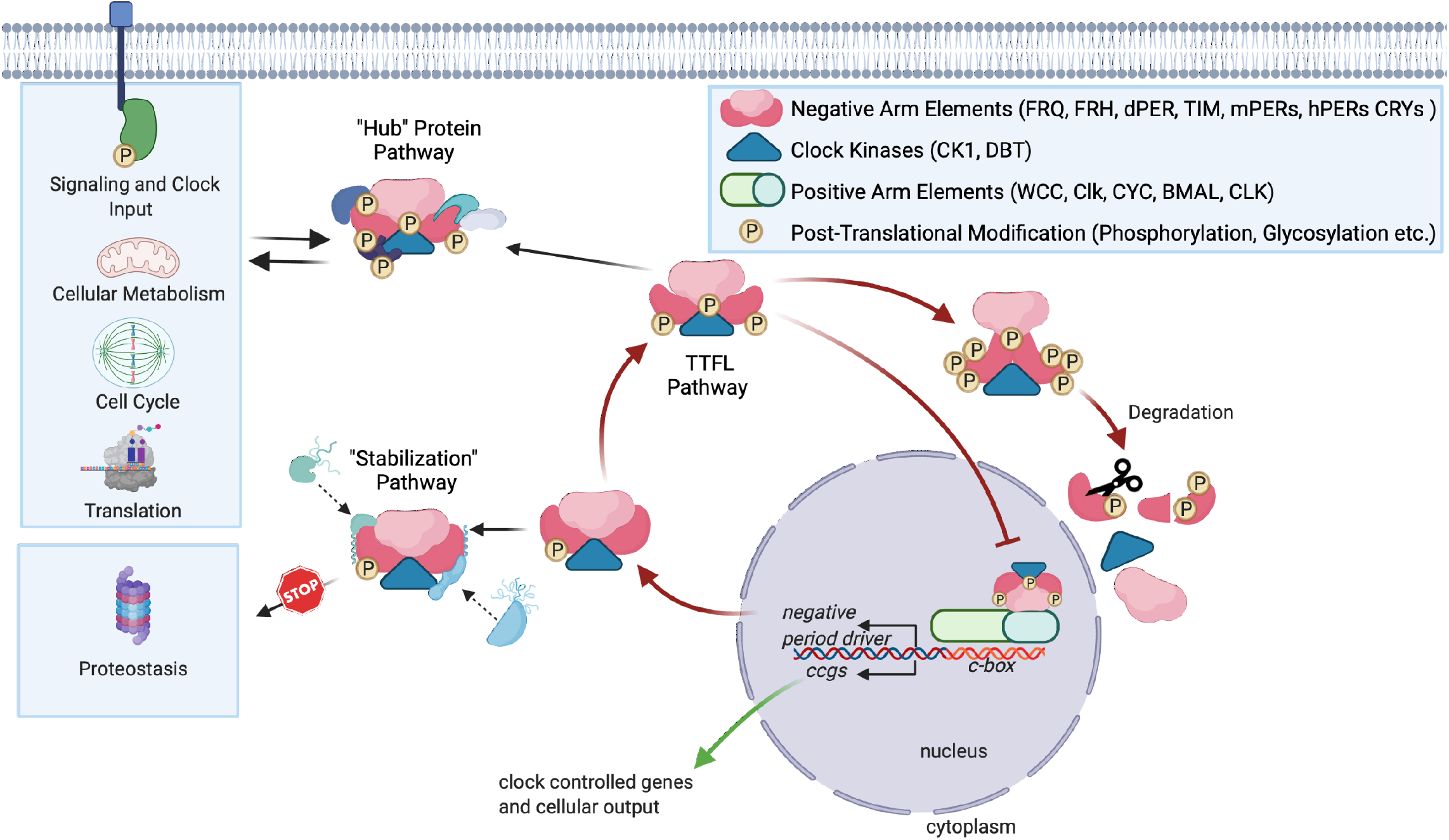
Clock negative arm proteins may act as a source of post-transcriptional clock regulation by acting as temporal “Hub” proteins and/or “proteosasis” regulators. Schematic representation of the classical circadian TTFL and the mechanisms by which we propose methods of negative arm post-transcriptional regulation occur. Proteins that bind negative arm macromolecular complexes can be temporally stabilized or phosphorylated, leading to downstream circadian post-transcriptional regulation in a variety of cellular pathways.

Many of the identified interactors regulate downstream pathways that are known to be under circadian post-transcriptional regulation and can themselves be regulated by phosphorylation ^19^. For example,14-3-3 zeta, a regulator of the c-Raf pathway, has been shown to be phosphorylated by CK1 ^92,93^. Further, the phosphorylation of CDC-28 is known to be one of the synchronizing links between the circadian and cell cycles, and the timing of its interaction with FRQ could facilitate temporal specificity of this transition between cell cycle phases ^71^. Therefore, our work suggests that the negative arm may act as a “Hub” protein to concentrate protein-protein interactions temporally to impart phosphorylation events on interactors in a circadian manner. This temporal “Hub” model, and the spatiotemporal specificity conferred by DBT (Figure 3), aligns with the suggested dPER foci heterogeneity speculated by Xiao et. al. ^94^.

Alternatively, or perhaps in parallel, our data also suggests a mechanism by which the negative arm temporally stabilizes interacting IDPs to impart post transcriptional regulation. While our predictions of protein disorder among interactors showed that IDPs were underrepresented in our dataset, many proteins that were IDPs interacted with FRQ at the nadir of their expression, suggesting that the negative arm may stabilize the protein to impart post-transcriptional regulation (Figure 5). This correlated with recent data showing the importance of the clock in proteostasis^81,82^. Importantly, our two models of negative arm post-transcriptional regulation, the “Hub” model and the “stabilization” model (Figure 6), are not mutually exclusive and do not necessitate a direct interaction between the negative arm and its cognate interactors, simply that a negative arm-regulated interaction occurred in the correct time and space.

Regardless of the mechanism by which the negative arm imparts post-transcriptional regulation, FRQ, dPER, and hPER have extensive interactomes, with functions that range across diverse gene ontologies (Figure 4). This aligns with the large and variable interactomes that are a hallmark of negative-arm proteins throughout eukaryotic circadian clocks ^23,27,31,95-98^. In FRQ, we found that 58 interactors, more than half of which (33) were IDPs, have rhythmic proteins without corresponding rhythmic mRNA. These 58 proteins accounted for ∼11% of all post-transcriptional regulation (rhythmic protein/mRNA discrepancies) reported in Hurley et al. 2018 ^19^. Further, many other FRQ interactors are responsible for the regulation of proteins under circadian post-transcriptional control ^19^. As proteins co-IPed from protein lysates have an intrinsic bias for soluble proteins, negative arm-membrane protein interactions may have been lost in our analysis and the negative arm-interactome may be larger than we report. In total, this suggests that negative arm-based post-transcriptional regulation could have great physiological reach and future research in the lab will focus on the validation and effect of this regulation.

## ACKNOWLEDGEMENTS

Figure 6 was created with BioRender.com. We thank Joshua Thomas and Catherine Mann for their assistance with creating the data processing script. We also thank Morgan Rhodes for her assistance with data formatting. We thank the Hurley lab members for their suggestions and support. This work was supported by an NIH-National Institute of Biomedical Imaging and Bioengineering Grant U01EB022546 (to J.M.H), an NIH-National Institute of General Medical Sciences Grant R35GM128687 (to J.M.H.), an NSF CAREER Award 2045674 (to J.M.H.), Rensselaer Polytechnic Startup funds (to J.M.H.), a gift from the Warren Alpert Foundation (to J.M.H.), and NIH-National Institute of Diabetes and Digestive and Kidney Diseases Grant R01DK124068 (to J.C.C.)

## AUTHOR CONTRIBUTIONS

Conceptualization, J.F.P, A.E.M, J.C.C and J.M.H.; Methodology, J.F.P, A.E.M, J.C.C and J.M.H; Investigation, J.F.P, A.E.M and L.S.B., Formal Analysis, J.F.P, A.E.M, C.L.K, S.C.A and W.B.F., Writing-Original Draft, J.F.P, A.E.M, Writing Review & Editing, J.F.P, A.E.M, J.C.C and J.M.H., Visualization J.F.P, A.E.M, C.L.K, S.C.A and W.B.F., Funding Acquisition, J.C.C. and J.M.H.

## METHODS

### Neurospora crassa strains, plasmids, and reagents

Four strains of *N. crassa* were used in this report (Supplemental Dataset 2). The cassette for strain WT8-1 was assembled as described in Colot et al. 2006, and transformed into an *frq* null (Δ*frq, bd*^*+*^, mat a) background at the *cyclosporin* locus to create the genotype *csr-1::frq*^*v5his103flag*^. Strain 1500-1 was created by transformative recombination at the native *frq* locus to fuse a V5-10His-3FLAG tag with an *hph* cassette to the C-terminal end of the protein FRQ (*frq*^V5H103FLAG^*::hph+, mus52::bar*). Transformations were performed as described ^101,102^ and screened by PCR. The negative control strain for our mass spectrometric analysis was Ku70a. Strains are listed in Supplemental Dataset 1. Race tube analysis of clock period was performed as described with slight modifications ^12,103-105^. For strain WT8-1, race tubes were filled with 15mL of Race Tube Media (1X Vogel’s, 1.5% Agar, 0.05% Glucose, 0.17% L-Arginine, and Biotin 0.005mg/mL), with cotton at the ends. For strains 328-4, 122, Ku-70a, and 1500-1, the same race tube media was prepared lacking glucose to facilitate conidial banding of non-banding strains. Tubes were inoculated at one end with conidia from slants grown on minimal media (see Fungal Genetics Stock Center website for recipe; 4-7 days of growth). Race tubes were grown in constant light (LL) at 25°C for 48 hours before being moved to 25°C constant darkness (DD) and marked at their growth front every 24 hours. WT8-1 tubes were analyzed for period using ChronOSX and all other strains were analyzed by hand.

To obtain tissue for NS-LC-MS/MS analysis, conidia was grown on minimal media slants and then harvested by adding 50 mL of liquid culture media (LCM) and vortexing (see Fungal Genetics Stock Center website for recipe) ^103^. Harvested conidia were resuspended in 4L of LCM. 4L cultures of *N. crassa* were grown at 25°C in LL before being moved to constant dark conditions (DD) and sampled at different circadian times (Supplemental Figure 2B). To obtain tissue for CRAFTY, conidia was grown on minimal media slants and harvested in 4-7 days by adding 2 mL of LCM and vortexing. Harvested conidia were added to 20 mL of LCM in a petri dish and left to grow at 25°C for one day in LL to form a mycelial mat. Plugs were cut from this mycelial mat and inoculated in 50 mL of LCM. *N. crassa* plugs were grown at 25°C in LL before being moved to constant dark conditions (DD) and sampled at different circadian times.

### Analysis of FRQ conform-ome using CRAFTY

CRAFTY assay and analysis was performed as described in Pelham et al. 2018 ^30^. Briefly, after the sample was grown for the allotted time, the culture was vacuum filtered and flash frozen, after which proteins were extracted using Protein Lysis Buffer with Halt™ Protease Inhibitor Cocktail diluted to a 1X concentration (Thermo Scientific™ 78430) ^30^. Protein concentration was measured by a Bradford Assay and standardized to 5 mg/mL at 4 °C. Thermolysin (8.9 uM) was added to the samples and the samples were aliquoted to a thermocycler. Samples were heated to 25 °C and at 30 second intervals the reaction was quenched using EDTA (20 mM) and SDS-PAGE loading buffer. Samples were analyzed using western blots, with 18.75ug of protein per lane in a 4-12% NuPAGE Bis-Tris gel. The primary antibody against the V5 tag on FRQ was sourced from Invitrogen (46-1157) and used at a 1:5000 dilution. The secondary antibody was Goat Anti-mouse IgG HRP conjugate sourced from Invitrogen (Invitrogen 313430) and used at a 1:25,000 dilution. The digestion rate was quantified and converted into protein half-life as in ^29^.

### Protein extraction and purification for mass spectrometric analyses of Neurospora

*Neurospora* tissue was harvested at the specified circadian times by vacuum filtration and proteins were extracted using Protein Lysis Buffer with Halt™ Protease Inhibitor Cocktail diluted to a 1X concentration (Thermo Scientific™ 78430) ^30^. Protein concentration was measured through Bradford Assay and standardized to 35 mg/mL. 45 mL of total protein was transferred to washed/charged Ni-NTA agarose beads (Invitrogen™ R90110) and incubated for 1 hour at 4°C with rotation before being washed. Proteins were eluted using Ni-NTA recommended elution buffer. All buffers were prepared as described in the manufacturer’s instructions. The Ni-NTA eluted proteins were incubated with washed/pre-conjugated anti-FLAG magnetic beads (Millipore™ M8823) overnight, washed, and eluted using Laemmle buffer. Beta-mercaptoethanol (BME) was added to an aliquot of protein elution for western blot analysis.

### Alkylation of Cysteine Residues

Proteins eluted from the anti-FLAG beads were reduced by adding 5mM of BME and incubated at 70 °C for 20 minutes. Iodoacetamide was added to the sample to a final concentration of 13mM and incubated at room temperature in the dark for 1 hour. The alkylation was quenched by adding an additional 5mM of BME at 25 °C for 15 minutes.

### Trichloroacetic Acid Precipitation and Protein Digestion

Alkylated protein products were mixed with cold 100% acetone and 100% trichloroacetic acid in a 1:8:1 ratio. This mixture was precipitated at -20°C for 1 hour before centrifugation at 11,500 rpm for 15 minutes at 4°C. The supernatant was discarded, and the protein pellet washed with 1 mL of cold 100% acetone and centrifuged again at 11,500 rpm for 15 minutes at 4°C. The supernatant was again discarded, and the protein pellet was air dried. The protein pellet was resuspended in 190 ul of 50 mM ammonium bicarbonate and digested with 20 ul of 1 ug/ul trypsin gold (Promega V5280) at 37°C for 4 hours.

### Analysis of FRQ Interactors by Mass Spectrometry

Samples were sent to the Mass-spec center at the University of Texas at Austin for analysis. Protein identification was performed via NS-LC-MS/MS using a Dionex Ultimate 3000 RSLCnano UPLC coupled to a Thermo Orbitrap Fusion. Prior to HPLC separation, the peptides were desalted using Millipore U-C18 ZipTip Pipette Tips following the manufacturer’s protocol. A 2 cm long x 75 µm I.D. C18 trap column was followed by a 75 µm I.D. x 25 cm long analytical column packed with C18 3 µm material (Thermo Acclaim PepMap 100). Run-time was 1 hour. The FT-MS resolution was set to 120,000, and 3 sec cycle time MS/MS were acquired in ion trap mode. Raw data was processed using SEQUEST HT embedded in Proteome Discoverer. Scaffold (Proteome Software) was used for validation of peptide and protein identifications with filtering to achieve 99% protein confidence or a 1% FDR.

### Neurospora Interactome Data Analysis

To classify proteins as interactors (both in experimental and control conditions), protein thresholds in Scaffold were set at 99.9% and peptide thresholds were set at 95%, which are the minimum probability that a protein or peptide are identified in the spectra. Only proteins with ≥5% coverage or ≥4 unique peptides were considered as “identified” in our dataset. All proteins identified in the Ku70 strain at all time points were pooled together and these proteins were removed from the list of proteins detected at any 1500-1 time point. The remaining proteins for each circadian time point in their respective grouping were compared and only proteins identified in at least two out of three of the triplicates were retained as valid FRQ interactors (Supplemental Figure 2E).

To create the networking script, we used the experimental data derived from our MS/MS data to define the primary interactions and used the STRINGdb database to define the secondary interactions. These primary and secondary interactions were mapped to NCU identifiers from their STRINGdb ID using FungiDB and Uniprot. The GO analysis used the NCU identifiers as inputs and outputs to associate GO terms with each gene. The R package “iGraph” was used to graph the network using the Fruchterman-Reingold layout and each node was color coded by the GO terms associated with it in a pie chart style. The “vertex.size” variable in the plotting function was changed to match the number of total nodes to limit overlapping.

Transcriptomics and proteomics datasets were taken from the 48-hour time series of *Neurospora crassa* from Hurley et al., 2018 ^19^. The imputed datasets were assessed for rhythmic gene expression or protein levels using the ECHO program ^106^. Rhythms in gene expression were classified as significant if ECHO calculated the absolute value of the amplitude change coefficient to be less than 0.15, the Benjamini-Hochberg-adjusted p value less than 0.05, and the period between 20 and 28 hours. Cell cycle genes and pathway information were taken from the KEGG (Kyoto Encyclopedia of Genes and Genomes) “Cell cycle - yeast – *Neurospora crassa*” pathway. The fitted dataset generated by ECHO from the triplicate time series was used for graphing and analysis. Peak expression times were the average of time points in CT corresponding to local maximums. Expression range was the difference between the average peak value and average trough value.

### Neurospora Interactome Western blots

FRQ, known FRQ interactors, and proteins of interest were verified through western blot analyses using the final purified protein elution from the FLAG pull down using a standard western blot protocol ^29^. Anti-V5 antibody (Invitrogen) was diluted 1:5,000. Anti-FLAG antibody was diluted 1:2,500 (F3165, Sigma). Immunodepleted anti-FRQ antibody was diluted 1:5000 ^103^. SuperSignal West Femto ECL (Pierce) was used for signal development. In order to verify that the purification process maintained FRQ’s core interactions, we performed western blot analysis using anti-FRH ^107^ (diluted 1:10,000), anti-WC1 ^108^ (diluted 1:250), and anti-WC2 ^109^ (diluted 1:250).

### Fly entrainment, collection, protein extraction, and affinity purification

Flies expressing 3XFLAG-tagged dPER(WT), or dPER(D755-809), were entrained in a 12 hr light/ 12 hr dark (12:12 LD) cycle for 3 days at 25°C and collected on the fourth day at indicated time-points by flash freezing on dry ice. Fly heads and bodies were separated using metal sieves and heads were used for protein extraction. Roughly 3 ml of heads were used per extraction. Fly heads were pulverized in liquid nitrogen and ground into fine powder using mortar and pestle. Powder was homogenized in lysis buffer as described above in (20 mM Hepes pH 7.4, 1 mM DTT, 0.1% Triton X-100, 0.5 mM phenylmethylsulfonyl fluoride (PMSF), supplemented with Complete EDTA-free Protease inhibitor cocktail (Roche Life Sciences)) by using a 40 ml loose dounce homogenizer (Wheaton). The homogenate was filtered through 70um cell strainers (Corning Life Sciences, Tewsbury, MA) to remove fly debris before centrifugation to separate nuclei from cytoplasmic fraction. Lysed cells were centrifuged at 4^0^C, 2,000 rpm for 20 minutes. Supernatant was collected as cytoplasmic fraction and supplemented to get final Affinity Purification (AP) buffer composition of 20 mM Hepes pH 7.4, 1 mM DTT, 0.1% Triton X-100, 150 mM NaCl, 10% glycerol, 1 mM MgCl_2_, 25 mM NaF, 0.5 mM EDTA, 0.5 mM PMSF, supplemented with Complete EDTA-free Protease inhibitor cocktail (Roche). Nuclear pellet was washed once with lysis buffer to eliminate any residual cytoplasmic extract. Proteins from nuclei were extracted with rotation for 1 hr in nuclear extraction buffer (20 mM Hepes pH 7.4, 10% glycerol, 350 mM NaCl, 0.1% Triton X-100, 1 mM DTT, 1 mM MgCl_2_, 0.5 mM EDTA, 25 mM NaF, 0.5 mM PMSF, supplemented with Complete EDTA-free Protease inhibitor cocktail (Roche)) at 4^0^C, along with the cytoplasmic fractions. 10 units of DNase I (Thermo Scientific, Waltham, MA) was added into the fractions during this one-hour incubation to minimize identification of non-specific protein interactions mediated by DNA-protein contacts. Post incubation, NaCl concentration for nuclear fraction was diluted down to 150 mM. Nuclear and cytoplasmic fractions were subsequently centrifuged at 27,000Xg for 15 minutes to remove cellular debris. Immunoprecipitation was performed at 4°C for 1 hr and 30 minutes with 200 μl α-FLAG M2 beads (Sigma, St. Louis, MO) followed by four 10-minute washes using AP buffer. Proteins were eluted twice with equal bead volume (200 μl) of AP buffer supplemented with 3XFLAG peptide (Sigma) at 200 μg/ml at room temperature for 15 minutes on a Nutator.

### In-gel tryptic digestion and liquid chromatography-tandem mass spectrometry (LC-MS/MS) for Drosophila samples

To identify proteins bound to dPER, elutions were concentrated in a 14% Tris-Tricine gel and excised as a single gel slice. Each gel band was subjected to in-gel reduction, alkylation, tryptic digestion and peptide extraction as described in Qian et al. (2008). Peptides were solubilized in 0.1% trifluoroacetic acid and analyzed by Nano LC-MS/MS (Dionex Ultimate 3000 RLSC nano System interfaced with a Velos-LTQ-Orbitrap (Thermo Fisher, San Jose, CA). Samples were loaded onto a self-packed 100 µm x 2 cm trap (Magic C18AQ, 5µm 200 Å, Michrom Bioresources, Inc.) and washed with Buffer A (0.2% formic acid) for 5 min with a flow rate of 10 µl/min. The trap was brought in-line with the analytical column (Magic C18AQ, 3 µm 200 Å, 75 µm x 50 cm) and peptides fractionated at 250 nl/min using a segmented linear gradient: 4-25% B (0.2% formic acid in acetonitrile) in 60min, 25-55% B in 30 min. Mass spectrometry data was acquired using a data-dependent acquisition procedure with a cyclic series of a full scan acquired in Orbitrap with resolution of 60,000 followed by MS/MS (acquired in the linear ion trap) of the 20 most intense ions with a repeat count of two and a dynamic exclusion duration of 30 sec.

### Analysis of dPER Interactors by Mass Spectrometry

For protein identification from MS raw data, peak lists in MASCOT generic format (MGF) were generated using Proteome Discover 1.3 (ThermoFisher). Data were searched against Uniprot database for *Drosophila melanogaster* proteins using a local version of the Global Proteome Machine (GPM) XE Manager version 2.2.1 (Beavis Informatics Ltd., Winnipeg, Canada) with X!Tandem version 10-12-01-1 to assign spectral data (Beavis 2006). Precursor ion mass error tolerance was set to ±10 ppm and fragment mass error tolerance to ±0.4 Da. Cysteine carbamidomethylation was set as a complete modification, methionine oxidation and deamidation at asparagine and glutamine residues were set as variable modifications. All LC-MS data were analyzed together in a MudPit analysis and individual data extracted to ensure that peptides that could be assigned to more than one protein were assigned consistently for all samples. The resultant identifications were filtered by log GPM expectancy score (log(e)) of -5 for protein and -1 for peptide.

Spectral counts were utilized to establish a cutoff value, to discriminate between false and true interactors of dPER. If a protein had spectral counts ≤4, in 2 out of 3 replicates, the protein was removed from the final interactome list. Any protein found to interact with dPER that had negative control spectral counts greater than or equal to 4, in 2 out of 3 replicates, was deemed a nonspecific interactor and was removed from all timepoints and fractionation. This approach to data processing filtered the spectral counts data to identify unique interactors for both wild type PER and PERΔ. After data processing, timepoint and sample type specific PPI lists were generated. To visualize the differences in PPIs between both time points and subcellular localization, Venn diagrams were created from the lists of interactors. The Venn diagrams were created in BioVenn ^110^ and jvenn ^99^. This approach for comparative analysis was conducted on both WT PER and PERΔ datasets.

### SLiM and Computational Analysis

The sequences of FRQ, dPER and hPER2 were run though the eukaryotic linear motif predictor (ELM) ^53^. The filter criteria were set for each organism, taxonomic cut offs were organism specific (FRQ as Fungi, dPER as *Drosophila melanogaster*, and hPER2 as *Homo sapiens*), and motif probability score of 100. Since the ELM database has a high rate of false positives, the SLiMs were verified and retained through their protein interactors by manually cross referencing with each interactome (FRQ with Dataset 1, dPER with Dataset 3 and hPER2 Oyama et al. 2019 ^21^). SLiMs were then plotted on a linear map of each protein (Figure 2). PTMs were plotted on each of the respective maps and curated from various sources (FRQ ^22^, dPER ^67^, and hPER2 ^111^). For the disorder plots FRQ, dPER and hPER2 sequences were run through the IUPred2A(L) tool which generated the disorder and ANCHOR prediction ^89^.

