## Supplemental Figures for "Conformational Changes in the Negative Arm of the Circadian Clock Correlate with Dynamic Interactomes Involved in Post-transcriptionally Regulated Processes"

Supplemental Table 1. Key for SLiM maps in Figure 2.

| Color on Map | SLiM Category | hPER2 | dPER | FRQ |
|---|---|---|---|---|
| | Calcineurin (PP2B) docking motif | x | x | x |
| | PP1 docking | | | x |
| | PP4 EVH1 docking motif | x | | |
| | Casein Kinase 1 phosphorylation site | x | x | x |
| | Casein Kinase 2 phosphorylation site | x | x | x |
| | GSK3 phosphorylation site | x | x | x |
| | CDK phosphorylation site | | | x |
| | PKA phosphorylation site | | | x |
| | LATS kinase phosphorylation motif | | x | |
| | N-glycosylation site | | | x |
| | Sumoylation site | x | | |
| | 14-3-3 binding phosphopeptide motif * | x | x | x |
| | Di-Arginine ER retention/retrieving signal | x | x | |
| | Autophagy related protein-LIR binding motif | x | | |
| | Clathrin box motif | x | | |
| | RIR binding motif on REV1 DNA polymerase | x | | |
| | Ciliary targeting motif | x | | |
| | C-terminal binding protein (CtBP) ligand motif | | x | |
| | eIF4E binding motif | | x | |
| | Phosphotyrosine ligands bound by SH2 domains | | x | |
| | SH3 ligand | | x | |
| | Ligand motif binding the CSL BTD | | x | |
| | N-Arginine dibasic convertase cleavage site | x | | |
| | Y-based sorting signal | x | x | x |
| | Endosome-Lysosome-Basolateral sorting signals | x | | |
| | Nuclear Export Signal | x | | |
| | Nuclear Localization Signal | x | | x |
| | MAPK docking | | | x |
| | SPOP SBC docking | | | x |
| | SCF ubiquitin ligase binding Phosphodegrons | | x | |
| | N-end Degron | x | | |
| | Phosphorylation Site | | | |
| | Glycosylation Site | | | |
| | Acetylation Site | | | |
| | Ubiquitination Site | | | |

Supplemental table 1. The color key for the SLiM maps in Figure 2.
Each SLiM category is represented by its corresponding color. An X in the hPER, dPER, or FRQ column indicate the presence of the SLiM on each respective map (Figure 2B, D and F).

Supplemental Figure 1. CRAFTY analysis of FRQ demonstrates temporally-specific conformations.

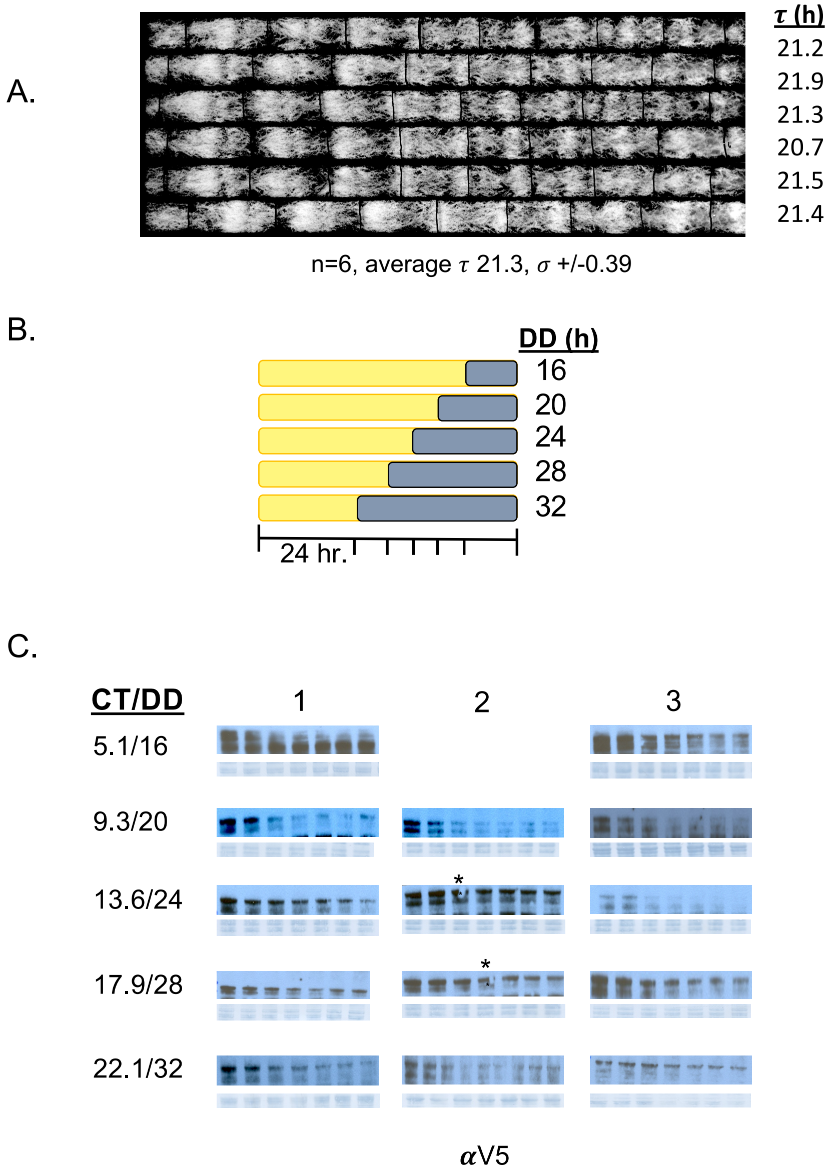


Supplemental Figure 1. CRAFTY analysis of FRQ demonstrates temporally-specific conformations. A.) Period determination of strain WT8-1 via race tube analysis. n= number of tubes analyzed, $\tau$=clock period in hours, $\sigma$=standard deviation. B.) Schematic of CRAFTY time course. Yellow=time in constant light (LL) and grey=time in constant dark (DD). C.) Western blots and corresponding amido black loading controls (below each blot) for each of the 3 replicates for the CRAFTY assay in Figure 1A (grey). DD 16 is an n of 2 as the second replicate was lost. * indicates lane was omitted from analysis due to blot irregularities.

Supplemental Figure 2. FRQ-centered macromolecular complexes are maintained in a tagged strain.

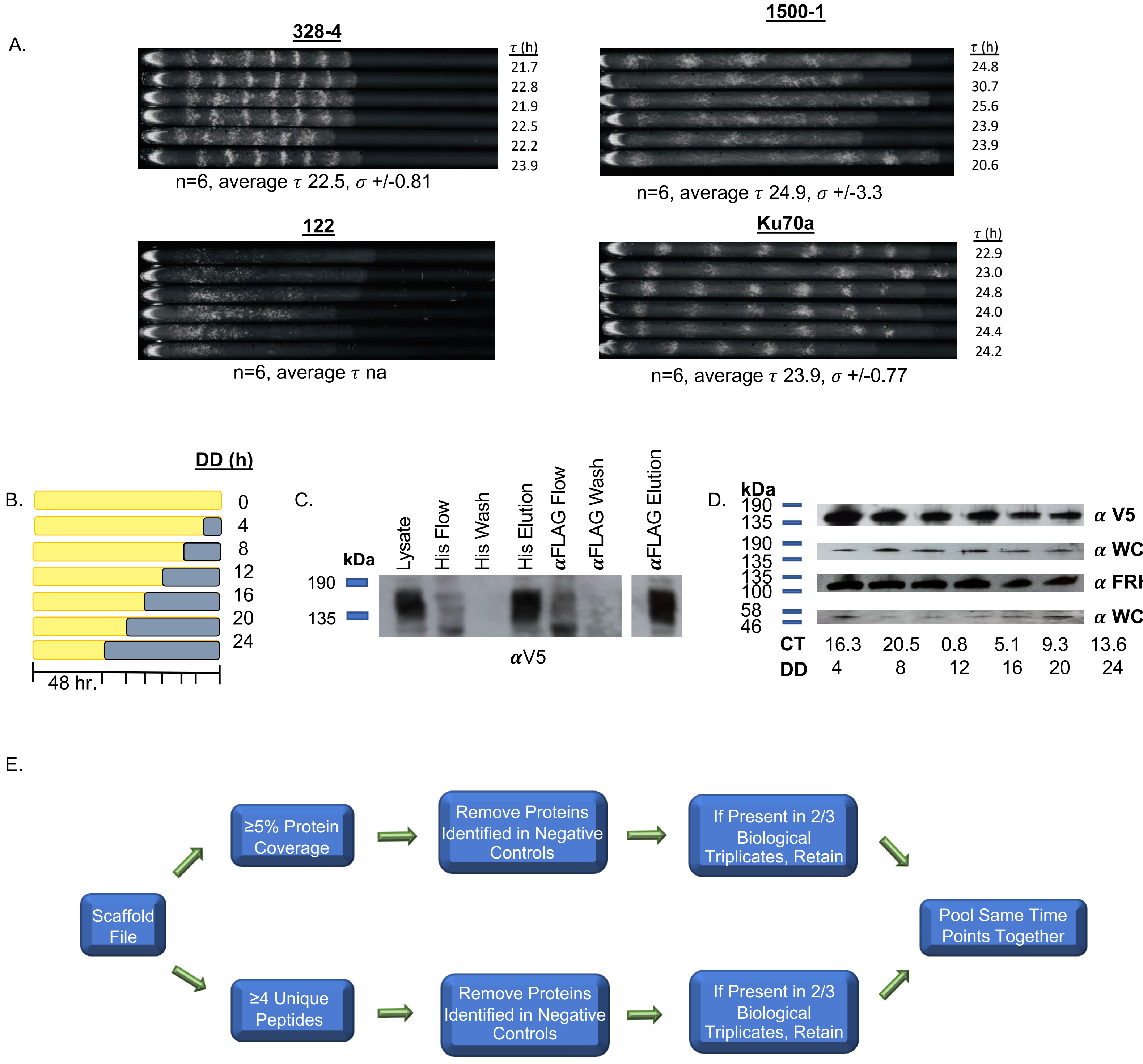


Supplemental Figure 2. FRQ-centered macromolecular complexes are maintained in tagged strain. A.) Period determination of strain 1500-1 via race tube analysis. n= number of tubes analyzed, τ=clock period in hours, σ=standard deviation. Race tube media lacking glucose was used to facilitate banding. B.) Schematic of NS-LC-MS/MS time course. Yellow=time in constant light (LL) and grey=time in constant dark (DD). C.) A western blot demonstrating the retention of FRQ at all purification stages. D.) Western blots of a co-IP using the 1500-1 strain with antibodies for the known FRQ interacting proteins (**α**-V5 for FRQ, **α**-FRH, **α**-WC-1, and **α**-WC-2). E.) A pictorial explanation of the LC-MS/MS data analysis method to define proteins identified in complex with FRQ.

Supplemental Figure 3. Proteins in negative arm macromolecular complexes are time specific and enriched for regions of disorder.

**A.**

| **All Time Points** | |
|---|---|
| FREQUENCY | NCU02265 |
| FRQ Interacting RNA Helicase | NCU03363 |
| Urease accessory protein ureG | NCU01511 |
| T-complex protein 1 subunit $\delta$ | NCU02839 |
| Glycerophosphocholine phosphodiesterase Gde1 | NCU01747 |

| **CT9.3- CT 0.8** | |
|---|---|
| WD repeat protein | NCU09162 |
| Protein kinase domain-containing protein | NCU05260 |
| DUF1649 domain-containing protein | NCU06174 |
| Nascent polypeptide-associated complex subunit alpha | NCU00635 |
| UPF0047 domain-containing protein | NCU09013 |

| **CT16.3-CT 9.3** | |
|---|---|
| nitrogen catabolic enzyme regulatory protein | NCU09068 |
| Telomere binding protein | NCU03830 |

| **CT16.3-CT 5.1** | |
|---|---|
| metalloprotease-15 | NCU05160 |

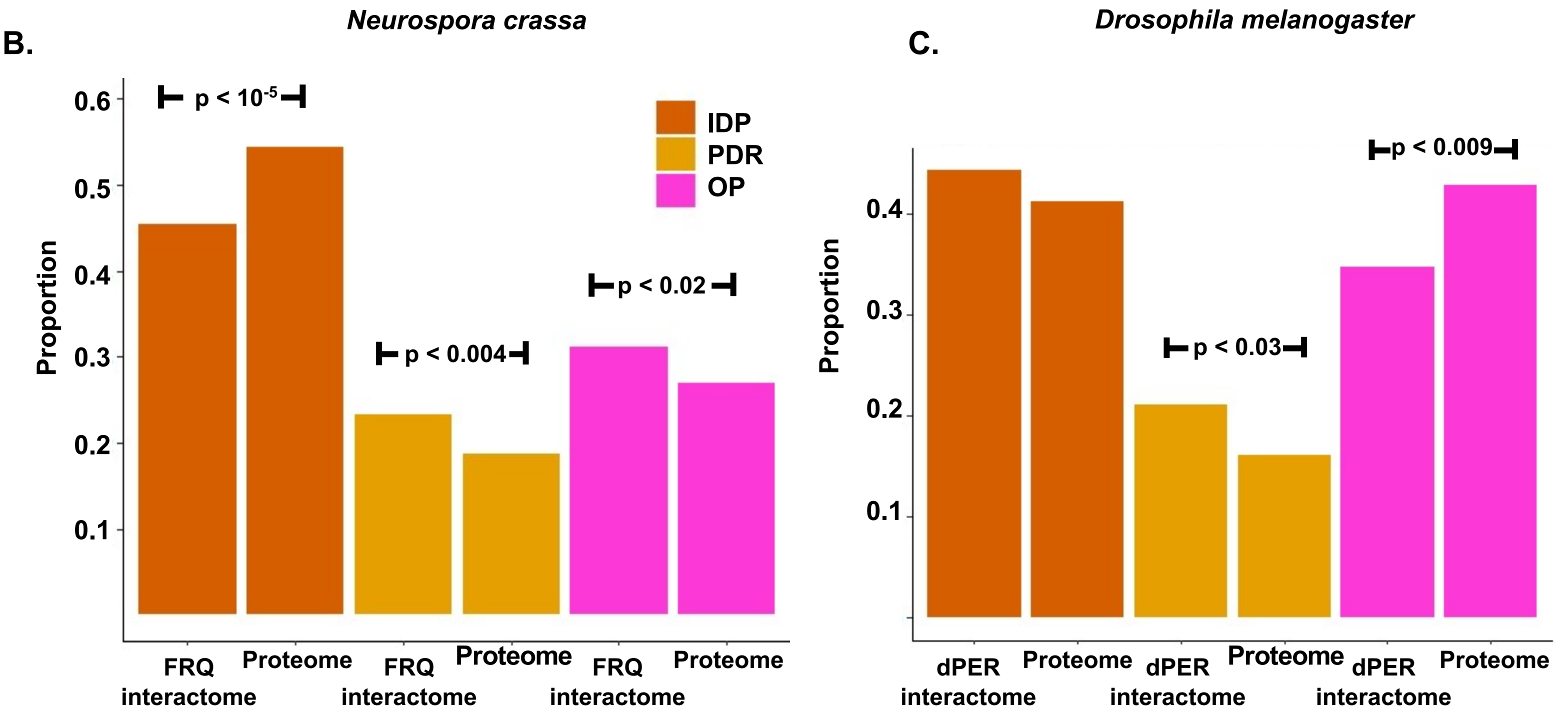


Supplemental Figure 3. Proteins in negative arm macromolecular complexes are time specific and enriched for regions of disorder. A.) List of proteins found in complex with FRQ across four or more contiguous time points. B.) A graphical representation of the proportion of proteins from each of the 3 classes of disorder found in the DD interactome plotted against the proportion of each of the 3 classes of disorder in the overall proteome of *Neurospora.* Fisher's exact test $p < .05$ indicates significance. C.) A graphical representation of the proportion of proteins from each of the 3 classes of disorder found in the dPER interactome plotted against the proportion of each of the 3 classes of disorder in the overall proteome of *Drosophila.* Fisher's exact test $p < .05$ indicates significance.

Supplemental Figure 4. Circadian oscillation of cell cycle-associated RNAs and proteins.

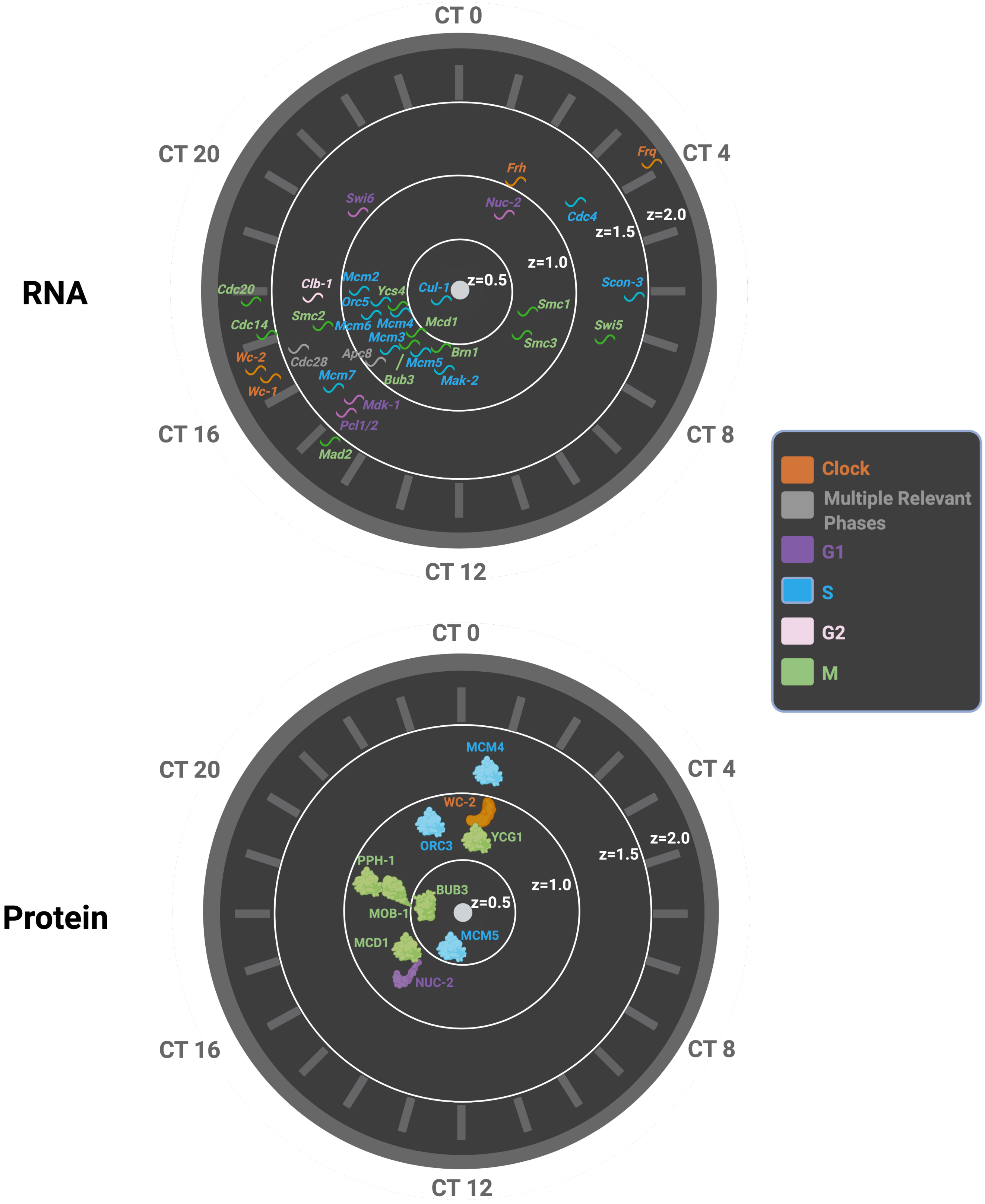


Supplemental Figure 4. Circadian oscillation of cell cycle-associated RNAs and proteins.
Rhythmic cell cycle-associated mRNA and proteins identified in Hurley et al. 2018 plotted on a clock style graph by the time of day of their peak and fold change of oscillation[19]. mRNA and protein are colored according to the stage in the cell cycle where they play a functional role.

Supplemental Figure 5. The composition of FRQ macromolecular complexes oscillated over the circadian day.

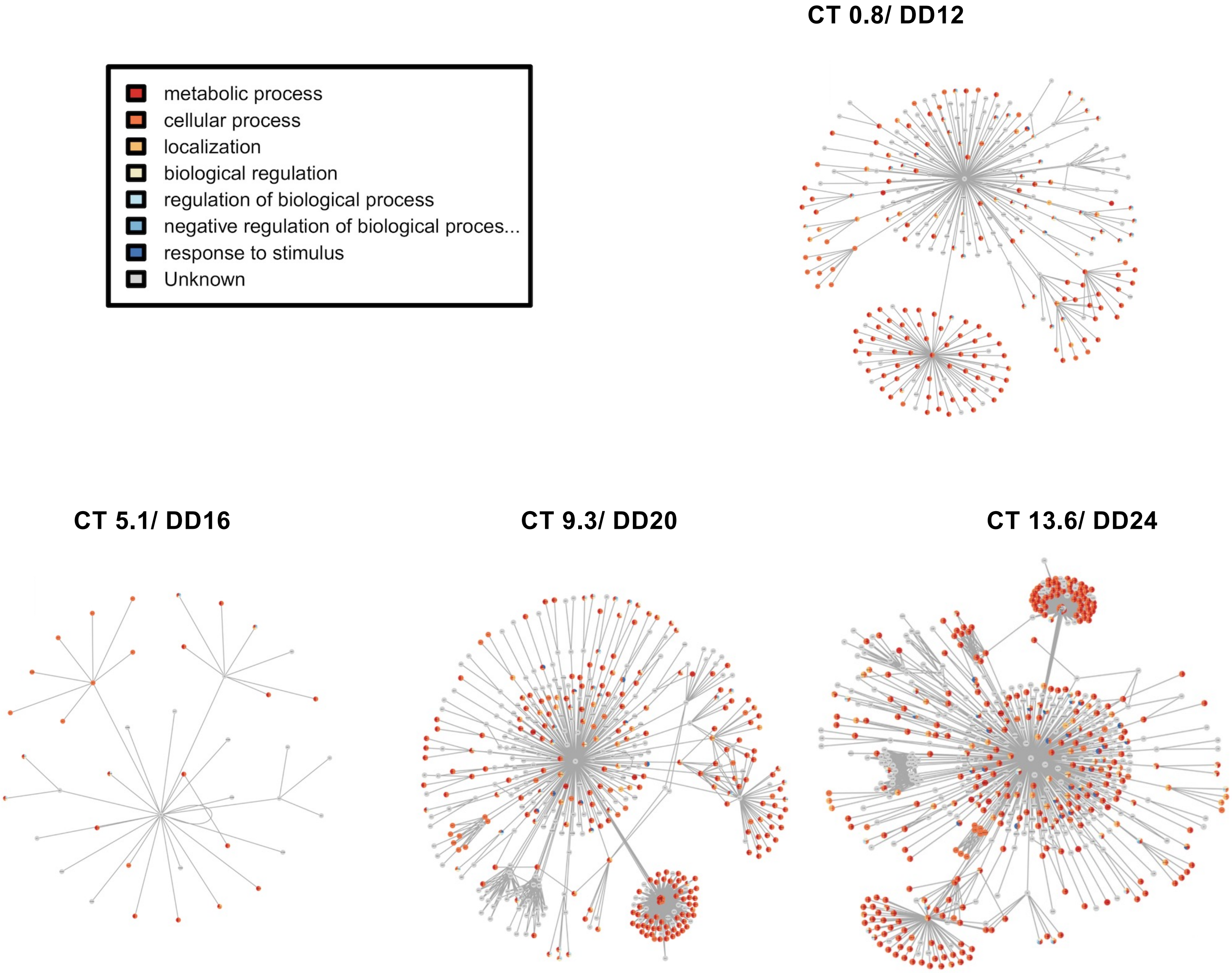


Supplemental Figure 5. The composition of FRQ macromolecular complexes oscillated over the circadian day. FRQ is in the center with radial lines connecting to interactors. Predicted secondary interactors are connected to their respective primary interactor. GO classifications are color coded according to the legend and proteins with multiple GO terms are shown by multiple slices of color in the biological processes category.

Supplemental Figure 6. The composition of dPER macromolecular complexes oscillates over the circadian day.

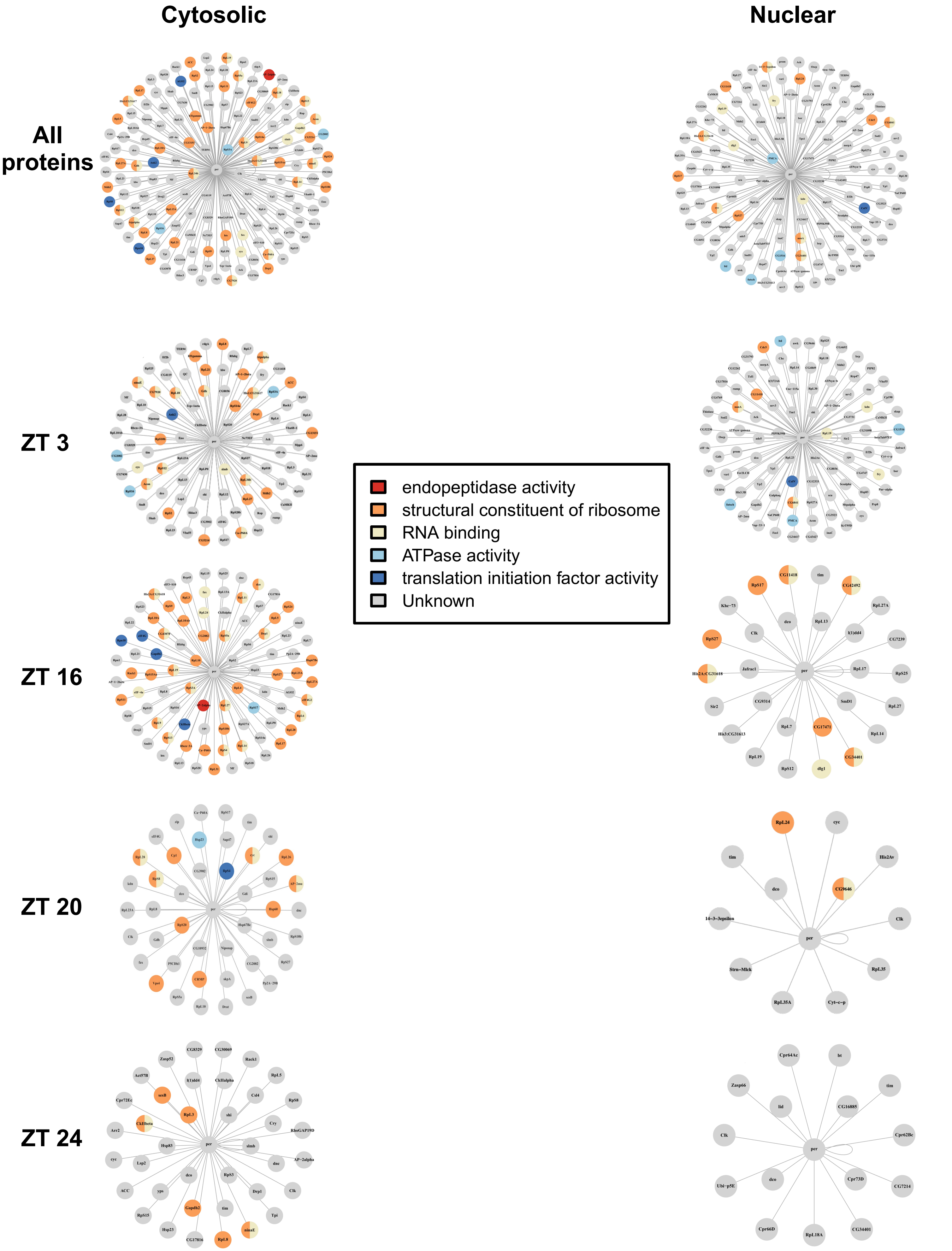

Supplemental Figure 6. The composition of dPER macromolecular complexes oscillates over the circadian day and by subcellular localization. dPER is in the center with radial lines connecting to primary interactors. GO classifications are color coded according to the legend and proteins with multiple GO terms are shown by multiple slices of color in the molecular function category.
